# Biodiversity-based innovations fortify agricultural growth in Asia-Pacific economies

**DOI:** 10.1101/845214

**Authors:** K.A.G. Wyckhuys, Y.H. Lu, W.W. Zhou, M.J.W. Cock, M.J. Furlong

## Abstract

The Green Revolution (GR) is widely credited with alleviating famine, mitigating poverty and driving aggregate economic growth over the past 6 decades. In Asia, GR technologies secured a tripling of rice output, with one-off germplasm improvements providing benefits beyond US$ 4.3 billion/year. Here, we unveil the magnitude and macro-economic relevance of parallel biodiversity-induced productivity growth in non-rice crops from 1918 to 2018. We empirically demonstrate how biological control (BC) defused invasive pest threats in multiple agricultural commodities, ensuring annually-accruing (on-farm) benefits of US$ 22.6 billion/year. Scientifically-guided BC permitted 73-100% yield-loss recovery in critical food, feed and fiber crops including cassava, banana, breadfruit and coconut. As such, BC promoted rural growth and prosperity even in marginal, poorly-endowed, non-rice environments. By placing agro-ecological innovations on equal footing with input-intensive measures, our work provides lessons for future efforts to mitigate invasive species, restore ecological resilience and sustainably raise output of global agri-food systems.

## Introduction

Global environmental change is undermining many of the UN Sustainable Development Goals, thus compromising endeavors to ensure economic growth, alleviate malnutrition, lift societal wellbeing and stabilize the Earth’s life-support systems (Griggs et al., 2013). Although agri-food systems sustain the livelihoods of >2.5 billion people and help meet dietary requirements of a swelling human population, they also contribute to global warming, ecosystem decay, and agro-chemical pollution (Tilman et al., 2001; Bernhardt et al., 2015; Maxwell et al., 2016; Springmann et al., 2018; Pretty et al., 2018). As such, agriculture-related habitat loss, natural resource degradation and a precipitous decline of biodiversity push the Earth system to a state that’s distinctly less conducive to societal development (Rockstrom et al., 2009; Potts et al., 2010). Given that the Asia-Pacific is responsible for roughly 50% of global agricultural production and its farm output has experienced a steady 3.8% annual growth over the last 5-6 decades (Alston & Pardey, 2014), the above externalities are believed to be particularly pronounced.

As a powerful engine of economic development (Johnston & Mellor, 1961), agriculture supplies food and raw materials and acts as a source of capital formation, residual labor, foreign exchange and purchasing power. Though often perceived as a stagnant ‘sunset’ industry, vigorous growth in smallholder agriculture permitted a rapid emergence and continual performance of Asia’s newly-industrialized ‘Tiger Cub’ economies (de Janvry, 2010; Berendsen et al., 2013). Post-1960s productivity growth in Asian agriculture has largely been credited to the Green Revolution (GR), i.e., one-off varietal improvements of cereal staples deployed under high-input ‘technology packages’ (Anderson et al., 1997; Pingali, 2012; Alston & Pardey, 2014). Founded on century-long progress in Mendelian genetics (Charnley, 2013), double-yielding, semi-dwarf rice varieties are heralded as key determinant of transformative growth in local agriculture. While improved rice germplasm surely unlocked the productive potential of large swathes of farmland, other biological innovations may have effected comparable technological change though their role has been obscured. Here, we aim to show how parallel achievements in insect biological control benefited Asia’s agricultural growth since the 1960s.

As a desirable, biodiversity-based agricultural innovation that conserves or enhances natural capital stocks (Sayer & Cassman, 2013), biological control (BC) is the scientifically-guided conservation, augmentation or introduction of beneficial organisms for the mitigation of (invasive or endemic) crop pests, pathogens and weeds. Over its 130-year history, biological control has achieved an effective suppression of at least 220 different invasive pestiferous insects worldwide, regularly at benefit-cost ratios of 5:1 to >1,000:1 (Naranjo et al., 2015; Heimpel & Cock, 2018; Heimpel & Mills, 2018). At those rates of return, 80% of documented benefits of the entire Consultative Group of International Agricultural Research (CGIAR) in certain geographies emanate from a narrow set of BC interventions (Maredia & Raitzer, 2010). At present, invasive pests inflict US$23–34 billion annual losses to Southeast Asian agriculture (Nghiem et al., 2013), slow rural economic growth and imperil food security in settings with fast-growing populations (Bradshaw et al., 2016; Savary et al., 2019). Being a self-propelling technology that requires limited stakeholder involvement, biological control is particularly suitable to alleviate invasive pest issues in remote settings and smallholder farming systems (Andrews et al., 1992).

While invasive species receive only scant attention by present-day scientists in the Asia-Pacific region (Peh, 2010), they were prime targets of thriving internationally-orchestrated development programs during the colonial period and latter half of the 20^th^ century (Waterhouse, 1998; Heimpel & Mills, 2018). BC interventions resolved pests such as the coconut scale *Aspidiotus destructor* Signoret or the banana skipper *Erionota thrax* (Linnaeus) that jeopardized economic prosperity, food security and welfare of entire nations (Cock, 2015; Spenneman, 2019). By restoring ecological balance and defusing a need for synthetic pesticides in invaded agri-food systems, BC preserved local biota and contributed to biodiversity conservation at large (Hoddle et al., 2004; Van Driesche et al., 2010; Hajek et al., 2016). Yet, following unintended negative consequences of a small number of unfortunate BC introductions made in the mid-1900s, regulatory hurdles and a more risk-averse attitude slowed BC progress and eclipsed its myriad societal benefits (Heimpel & Cock, 2018). Now, as pest invasions continue unabated (Wild, 2017) and environmental consequences of their improper mitigation become more apparent (Ricciardi et al., 2011; Bellard et al., 2017; Sanchez-Bayo & Wyckhuys, 2019), an *ex-post* valuation of BC economic benefits and its relative contribution to agricultural growth can shine new light on this valuable practice.

A ‘food systems’ perspective is well-suited to blend biophysical facets of global environmental change, i.e., species invasion, pest-induced crop loss or BC-mediated loss recovery, with socioeconomic parameters to optimally gauge their societal outcomes (Ericksen, 2008; Ingram, 2011; Hackmann et al., 2014; Liu et al., 2015). Using an integrative food systems lens, we a) systematically assess occurrence and fate of historical insect BC introductions across 26 different Asia-Pacific geopolitical entities, over a 1918-2018 timespan; b) quantify monetary impact of pest invasions and ensuing BC interventions at varying spatial scales, and c) contrast locality-specific productivity growth trajectories (i.e., resulting from progressive technological change; Ruttan, 2002) for rice and various non-rice commodities, reflective of either concurrent or disparate GR and BC interventions. Our work provides compelling evidence of how BC innovations yielded food system outcomes comparable to GR seed x fertilizer technologies, generating agricultural growth and prosperity for multiple Asia-Pacific nations including its ‘Tiger Cub’ economies.

## Results

### Invasion history & biological control efforts

Over the 1918-2018 timeframe, the BIOCAT database contained records of 160 insect natural enemies, employed against 77 invertebrate pests in agriculture, human health and livestock production sectors of 26 different Asia-Pacific geopolitical areas (Fig. 1). Natural enemies mainly comprised species within the taxonomic orders Hymenoptera (56%), Coleoptera (30%) and Diptera (11%), while BC targets largely included Hemiptera (38%) and Lepidoptera (25%) (Fig. 2). A total of 17, 21 and 25 different species of Encyrtidae, Braconidae and Coccinellidae were used for BC purposes, with most releases conducted in Fiji (18%), Northern Marianas (12%), Guam (11%), Indonesia (8%) and American Samoa (6%). Invasive pests of coconut were most commonly targeted followed by those on banana and citrus, with a respective 21, 14 and 12 different areas receiving (often multiple) BC releases on one of the three crops. Biological control was effectively used to mitigate livestock pests, i.e., buffalo fly *Lyperosia exigua* in Papua New Guinea (PNG), while the elephant mosquitoes *Toxorhynchites amboinensis* (Doleschall) en *T. splendens* (Wiedemann) were successfully employed against disease-carrying *Aedes* spp. and *Culex* spp. in at least five different areas.

**Figure 1.**
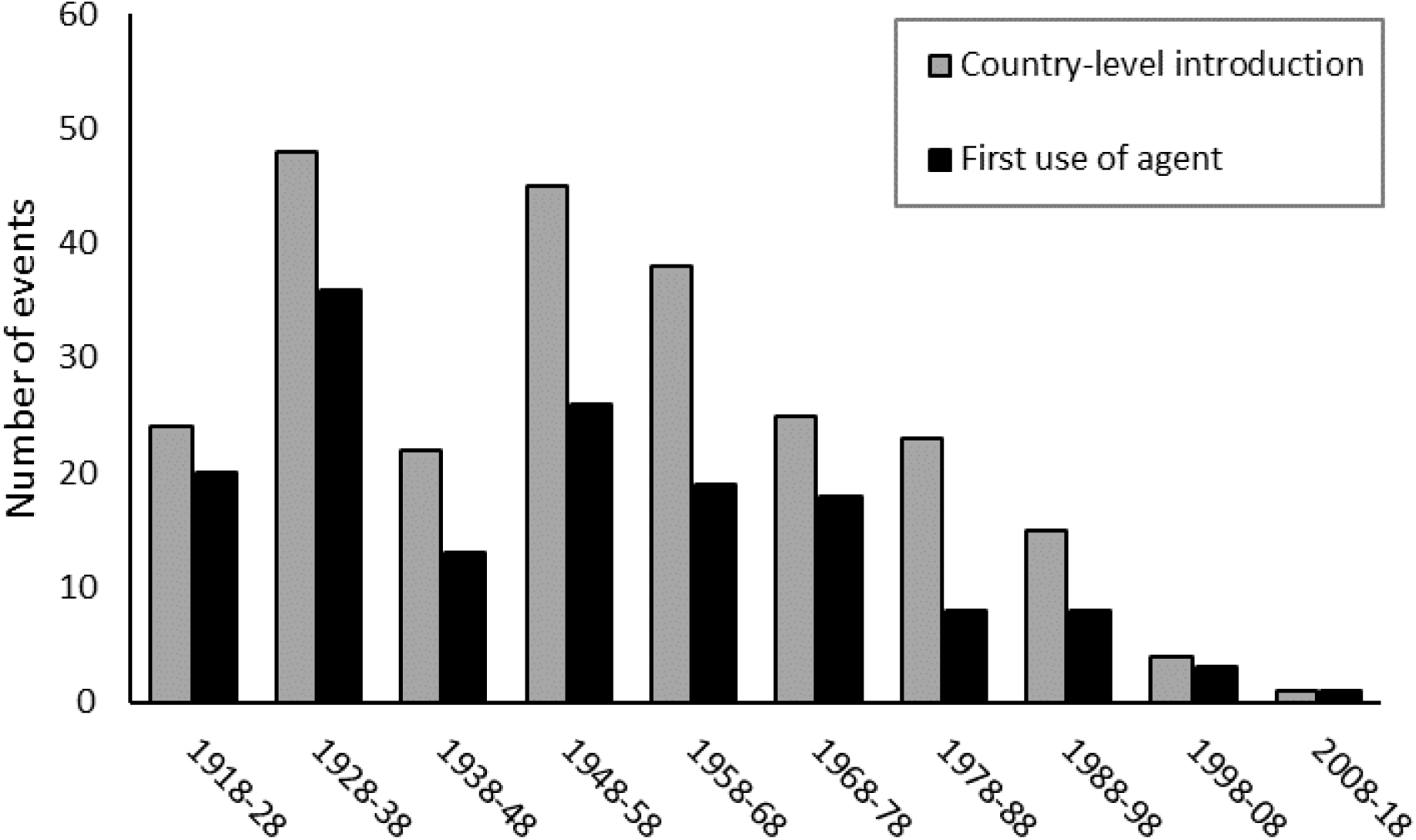
Timeline of insect biological control introductions across the Asia-Pacific region (26 countries). The total number of country-level introductions and first regional deployments of a given biological control agent is depicted for successive decades, over a 1918-2018 window. All introductions pertain to the use of insect natural enemies for insect pest management purposes in local agriculture, human health and livestock production sectors. Records are drawn from the BIOCAT database.

**Figure 2.**
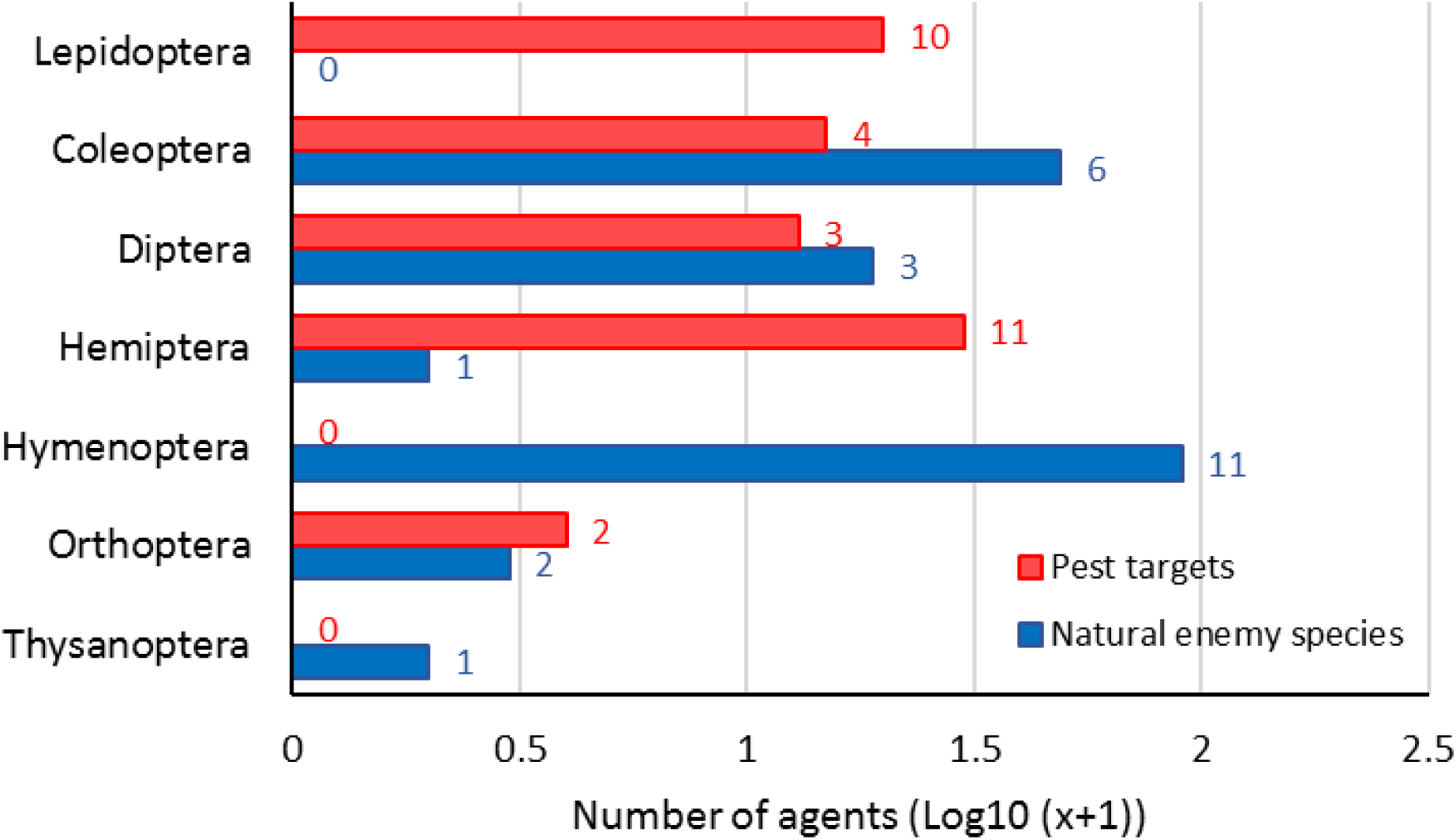
Insect natural enemies, and their related pest targets, as employed in historic biological control programs in the Asia-Pacific region. For each insect order, the number of agents (i.e., individual pest or natural enemy species) is listed. Next to each bar, the number of families within a given taxonomic category is indicated. Records cover all biological control interventions, including failed and partially/fully successful interventions, from 1918 onwards. Records are drawn from the BIOCAT database.

Out of 264 unique country-level BC interventions, 101 achieved establishment of the introduced BC agent (or so-called ‘natural enemy’) with an ensuing BC success rate of 38% (full or partial control). Once established, members of the Hymenoptera and Diptera attained BC success rates of 46% and 40%, respectively. Overall, 38% introduced species attained lasting establishment and 47% of those subsequently secured full or partial pest control within at least 1 geopolitical area. Expanding our analyses beyond BIOCAT database entries, we recorded a total of 76 BC agents that provided full or partially-successful control of 45 invasive pest targets in local agriculture and livestock production sectors. Non-exhaustive literature analysis thus yielded a total of 175 country-level BC interventions that resulted in (partially) successful pest control across the Asia-Pacific.

### Productivity shocks and producer impacts

Invasive pests caused important reductions in the primary productivity of a range of crops, e.g., the hispid *Brontispa longissima* (Gestro) inflicted 50-70% losses in coconut, the scale insect *A. destructor* regularly led to total loss of coconut, breadfruit and banana, while the fruit fly *Bactrocera dorsalis* (Hendel) impacted yield and marketability of mango by 82-95% (Supplementary Fig. 1). Using 2015 production and pricing statistics, historical pest invasions caused an aggregate farm-level economic loss of US$ 1.2 ± 3.8 billion per country annually (mean ± SD; *n*= 23 countries), with the highest impacts for Indonesia ($ 17.87 billion). Per crop, an annual aggregate economic impact of US$ 1.2 ± 2.2 billion was recorded, with the highest pest-related losses for coconut ($ 8.20 billion), cassava ($ 6.8 billion) and banana ($ 4.3 billion) (Fig. 3, 4). Following biological control, pest-induced losses were offset by an estimated 81%, for a total aggregate amount of US$ 22.62 billion annually. This represents 2.5 ± 3.8% of the current GDP (PPP) of pest-invaded countries, with the highest contribution to the national economy for Vanuatu (14.7%), Tonga (9.1%) and Solomon Islands (8.6%) (Table 1).

**Table 1.**
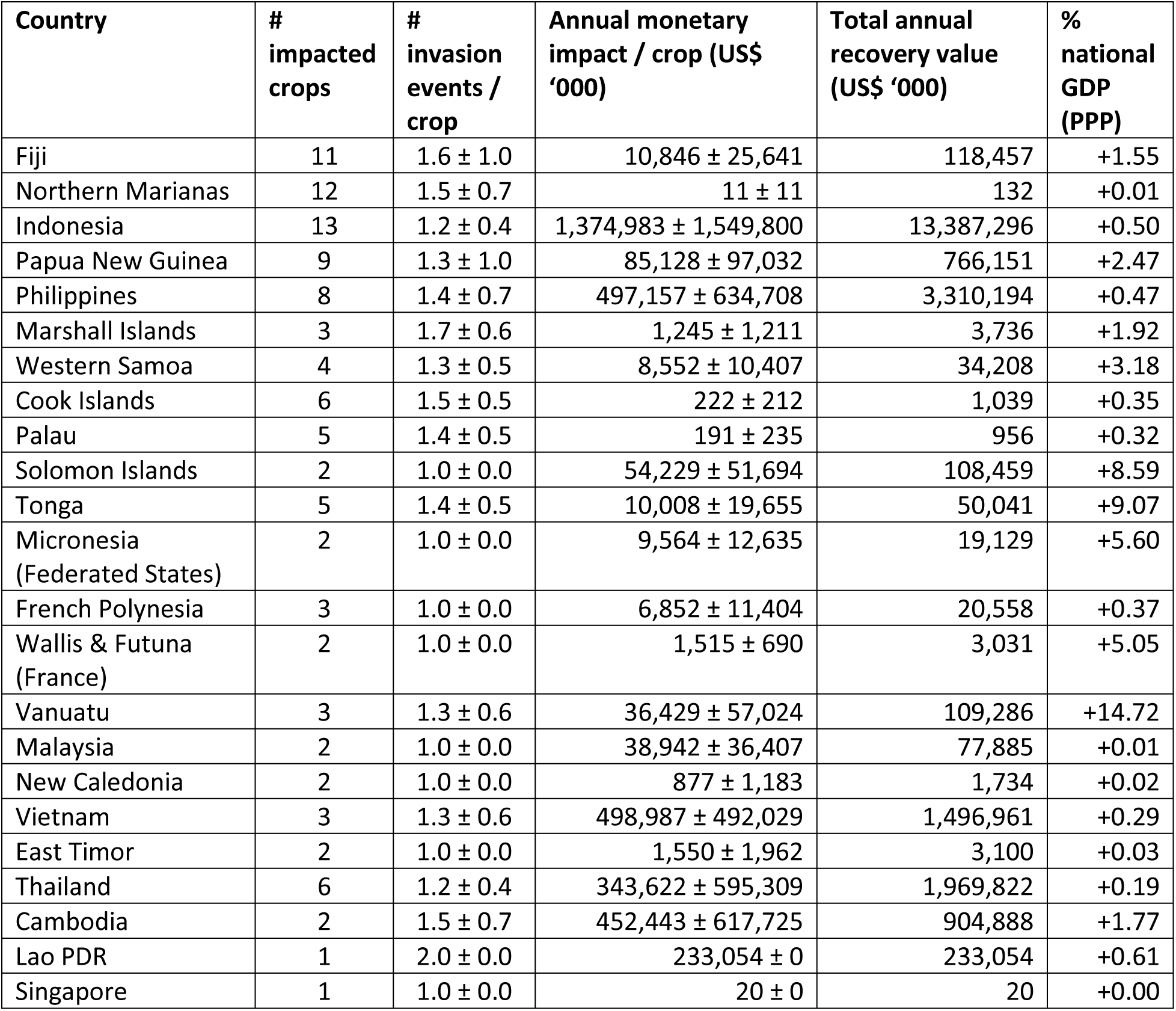
Country-level monetary impact of invasive insect pests and ensuing biological control interventions. Annual impact figures (mean ± SD) are exclusively computed for invasive insect pests that were either partially or fully controlled through biological control, over a 1918-2018 period. Patterns are shown for 23 countries in the broader Asia-Pacific region. Per country, aggregate recovery value for all biological control interventions is presented in absolute terms and as % of national GDP (PPP; 2015 US$). For crops subject to multiple historic pest invasions in a given country, only the biological control intervention with the highest recovery value is taken into consideration.

| Country | # impacted crops | # invasion events / crop | Annual monetary impact / crop (US\$ '000) | Total annual recovery value (US\$ '000) | % national GDP (PPP) |
| --- | --- | --- | --- | --- | --- |
| Fiji | 11 | 1.6 ± 1.0 | 10,846 ± 25,641 | 118,457 | +1.55 |
| Northern Marianas | 12 | 1.5 ± 0.7 | 11 ± 11 | 132 | +0.01 |
| Indonesia | 13 | 1.2 ± 0.4 | 1,374,983 ± 1,549,800 | 13,387,296 | +0.50 |
| Papua New Guinea | 9 | 1.3 ± 1.0 | 85,128 ± 97,032 | 766,151 | +2.47 |
| Philippines | 8 | 1.4 ± 0.7 | 497,157 ± 634,708 | 3,310,194 | +0.47 |
| Marshall Islands | 3 | 1.7 ± 0.6 | 1,245 ± 1,211 | 3,736 | +1.92 |
| Western Samoa | 4 | 1.3 ± 0.5 | 8,552 ± 10,407 | 34,208 | +3.18 |
| Cook Islands | 6 | 1.5 ± 0.5 | 222 ± 212 | 1,039 | +0.35 |
| Palau | 5 | 1.4 ± 0.5 | 191 ± 235 | 956 | +0.32 |
| Solomon Islands | 2 | 1.0 ± 0.0 | 54,229 ± 51,694 | 108,459 | +8.59 |
| Tonga | 5 | 1.4 ± 0.5 | 10,008 ± 19,655 | 50,041 | +9.07 |
| Micronesia (Federated States) | 2 | 1.0 ± 0.0 | 9,564 ± 12,635 | 19,129 | +5.60 |
| French Polynesia | 3 | 1.0 ± 0.0 | 6,852 ± 11,404 | 20,558 | +0.37 |
| Wallis & Futuna (France) | 2 | 1.0 ± 0.0 | 1,515 ± 690 | 3,031 | +5.05 |
| Vanuatu | 3 | 1.3 ± 0.6 | 36,429 ± 57,024 | 109,286 | +14.72 |
| Malaysia | 2 | 1.0 ± 0.0 | 38,942 ± 36,407 | 77,885 | +0.01 |
| New Caledonia | 2 | 1.0 ± 0.0 | 877 ± 1,183 | 1,734 | +0.02 |
| Vietnam | 3 | 1.3 ± 0.6 | 498,987 ± 492,029 | 1,496,961 | +0.29 |
| East Timor | 2 | 1.0 ± 0.0 | 1,550 ± 1,962 | 3,100 | +0.03 |
| Thailand | 6 | 1.2 ± 0.4 | 343,622 ± 595,309 | 1,969,822 | +0.19 |
| Cambodia | 2 | 1.5 ± 0.7 | 452,443 ± 617,725 | 904,888 | +1.77 |
| Lao PDR | 1 | 2.0 ± 0.0 | 233,054 ± 0 | 233,054 | +0.61 |
| Singapore | 1 | 1.0 ± 0.0 | 20 ± 0 | 20 | +0.00 |

**Figure 3a.**
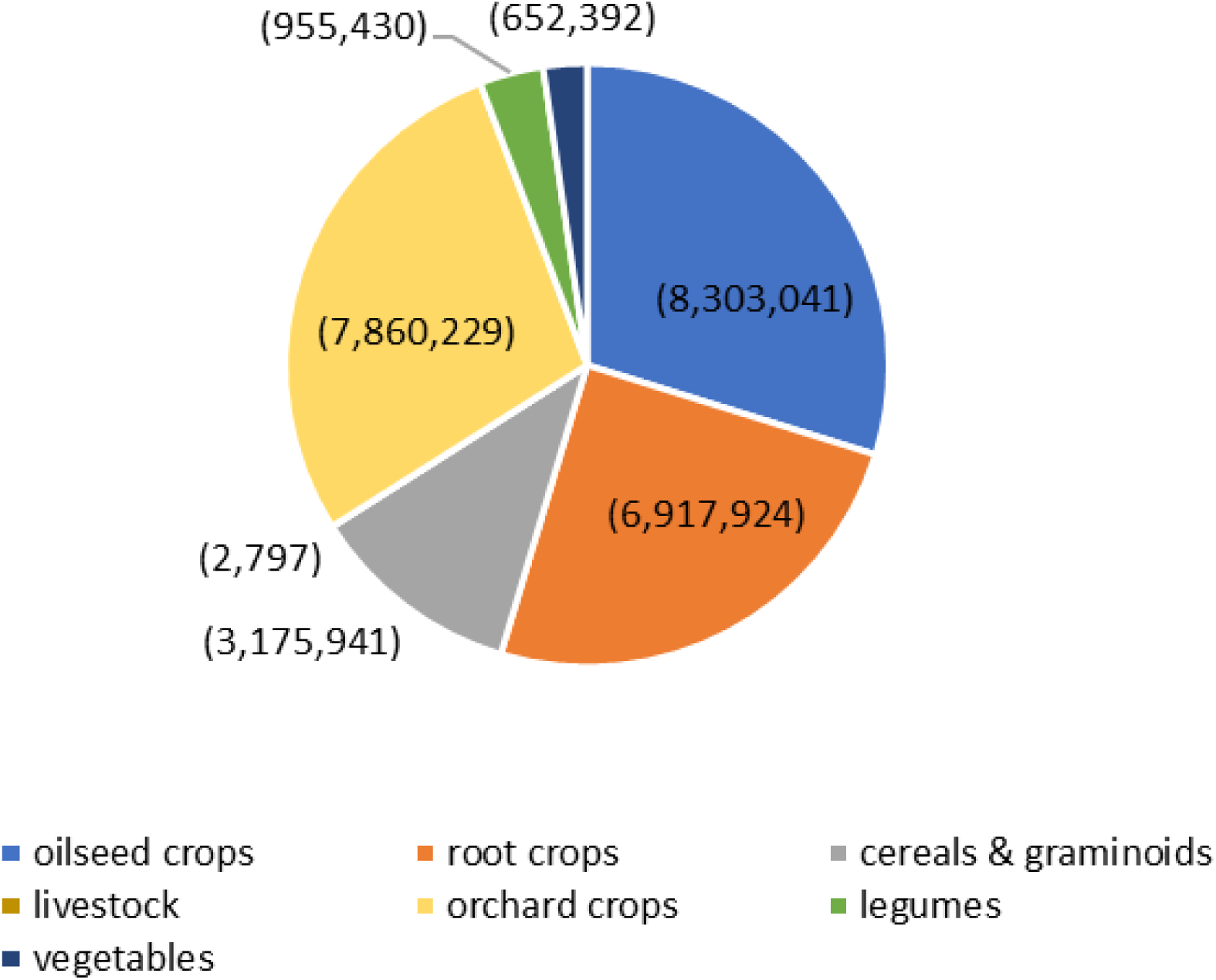
Aggregated annual economic impact of invasive pests (US$ ‘000), as partially or completely resolved through biological control during 1918-2018. The annual monetary impact of invasive agricultural pests amounts to US$ 27.87 billion (23 countries; 2015 US$), capturing historic invasion events in multiple cropping systems and livestock operations. Oilseed crops include coconut and oilpalm; root crops cover cassava and taro; while orchard crops cover banana, breadfruit, cocoa, coffee and a range of perennial or annual fruit crops. Records are drawn from the BIOCAT database and complemented with extensive literature revisions.

**Figure 3b.**
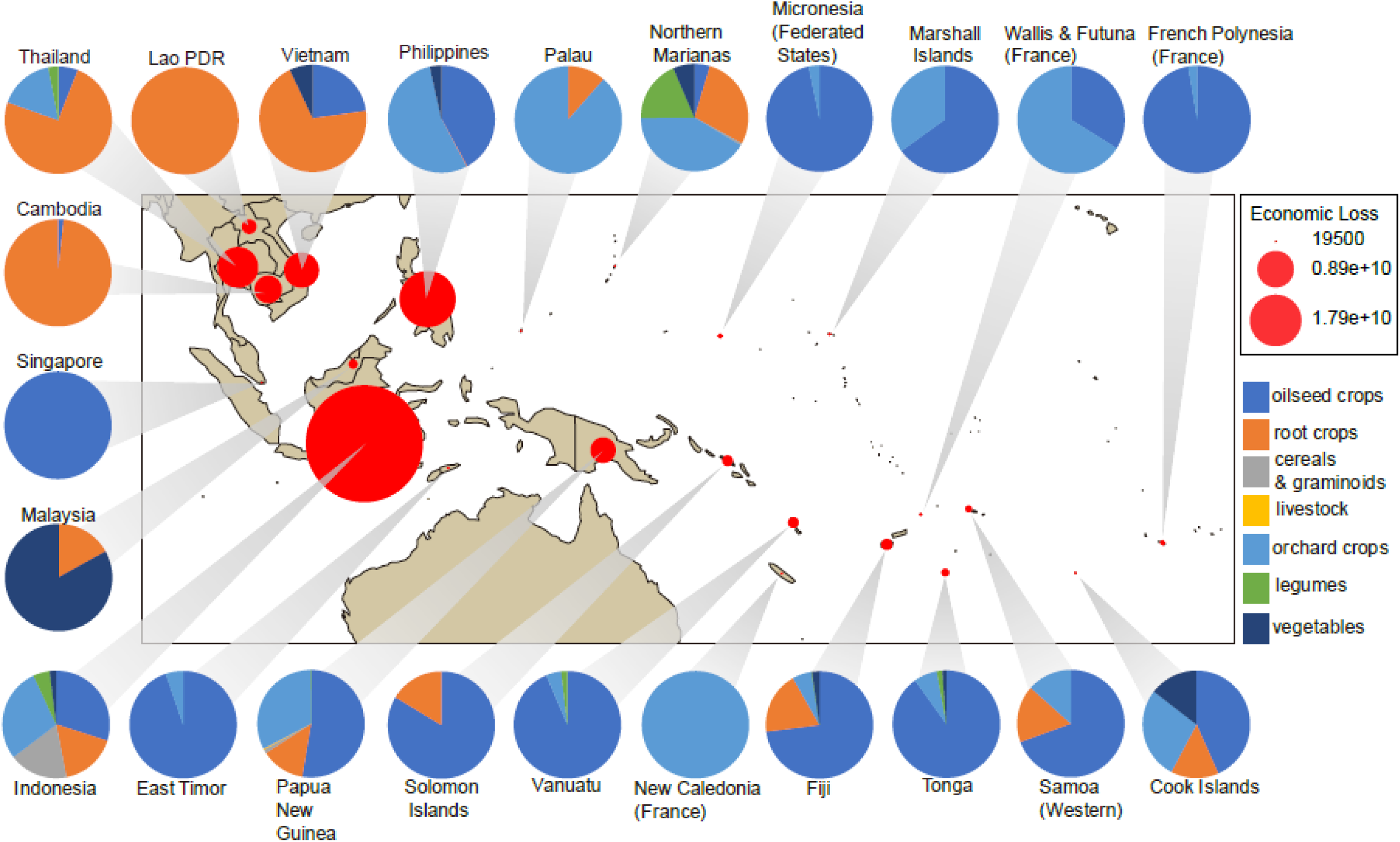
Country-level economic impact (US$) of historic pest invasions as either partially or completely resolved through biological control. For each country or geopolitical entity, the size of the red circle reflects the aggregate annual monetary impact of invasive crop and livestock pests over 1918-2018. Country-specific pie charts indicate the exact portfolio of commodities that benefited from invertebrate biological control, with the size of each slice reflecting the economic impact of respective pest invaders.

### Econometric analyses

Agricultural growth trajectories of the various countries differed substantially, with steep growth (i.e., intensification of labor use relative to land) in rice-cultivating economies such as Indonesia, Philippines, Thailand or Laos (Supplementary Fig. 2). In contrast, productivity paths of Melanesian countries were relatively flat (i.e., intensification of land use relative to labor) while growth was comparatively stagnant in e.g., Timor Leste, Tonga or Solomon Islands. Multiple countries that benefited from a succession of (land-saving) BC innovations (Table 1) thus experienced rapid productivity growth.

In 1961, all countries except Viet Nam possessed an agriculture sector dominated by non-staple commodities (in value; Fig. 4). For the six rice economies, aggregate value of the non-rice sub-sector was 1.3-22.9 times higher than that of paddy rice. For Pacific Island communities, value of the non-staple sub-sector was 4.1-25.7 times higher than that of taro and sweet potato combined.

**Figure 4.**
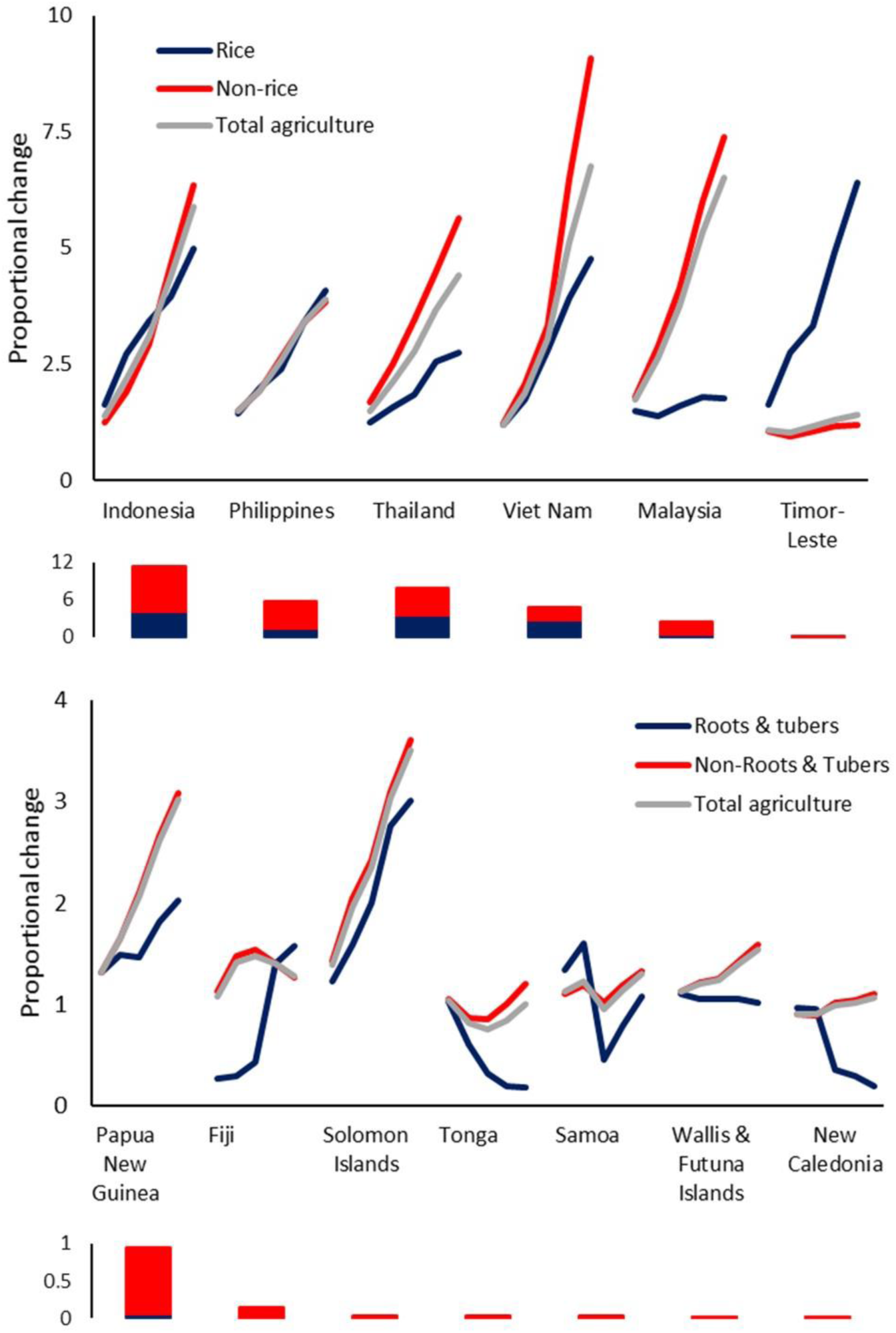
Differential growth trajectories of agricultural sub-sectors (i.e., food staple, non-staple crops) for selected geopolitical entities within the Asia-Pacific region, over 1961-2016. Patterns are presented for geopolitical entities that either rely upon rice or root & tuber crops (i.e., taro, sweet potato) as primary food staples. Within each graph, countries are ranked in order of a declining aggregate monetary value of biological control interventions. For either primary food staples (red color) or non-staple crops (blue color), interdecadal changes in their respective agricultural output are computed against a 1961-1970 baseline, as shown as a bar chart below each graph and expressed in $ billion PPP. Agricultural output is expressed in monetary terms, as total value of agricultural production (2004-2006 average PPP $; FAO 2019).

BC-reliant countries such as Philippines and Indonesia experienced comparatively rapid growth of the rice sub-sector over a respective two or three decades since the 1960s (Fig. 4), with the most vigorous rice productivity growth in Indonesia during the 1970s (Supplementary Fig. 3). All remaining rice economies were typified by superior growth of the non-rice subsector over >5 decades. For all Pacific Island communities except Samoa, initial pace of growth in the non-staple sub-sector surpassed that of key food staples (i.e., taro, sweet potato). Overall, aggregate agricultural growth in a specific geography mirrored that of the respective non-staple subsector.

Country-specific time paths revealed different sources of productivity growth for various BC-reliant commodities, as compared to paddy rice and other locally-relevant food staples (Fig. 5). Here, we attribute productivity growth in rice primarily to GR germplasm enhancement and high-input technology packages, while growth trajectories in non-staple crops are mediated to lesser extent by GR and can be partially ascribed to BC. In all rice economies, rice crops experienced substantial area increases in addition to max. 303-494% yield growth against a 1961 baseline. In BC-intensive countries such as Indonesia, max. yield growth in banana (1,222%), cassava (323%) and maize (572%) surpassed that of rice (303%) and can be partially ascribed to an effective recovery of pest-induced 81%, 73% and 25% yield losses in the respective crops. In rice economies, key BC-recipient crops such as coconut experienced yield growth between 126% (Philippines) and 279% (Vietnam). Furthermore, rice production areas in Melanesia contracted and local yield levels of several BC-recipient oilseed and orchard crops experienced modest to significant increases (i.e., coconut 145%, banana 148%, sugarcane 203%).

**Figure 5.**
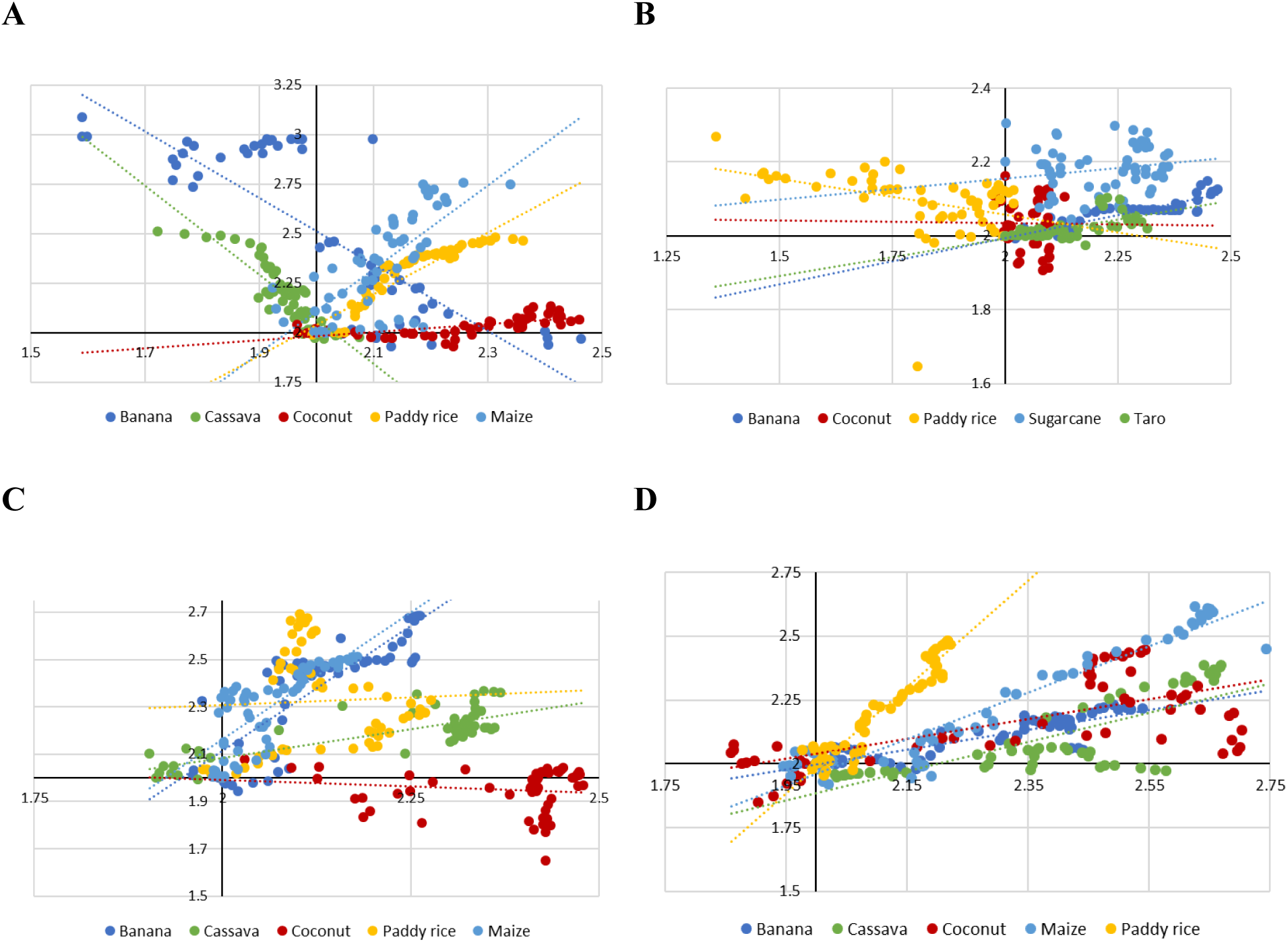
Time paths of productivity growth for different agricultural crops, over a 1961-2016 window. Inter-annual percentual growth (Log10) in area (Y axis) and yield (X axis) for different crops are measured against a 1961 base year. Trends are presented for Indonesia (A), Melanesia (B), Philippines (C) and Vietnam (D). High-impact biological control interventions are made in banana (Ind, Mel, Phi), cassava (Ind, Vie), sugarcane (Mel), coconut (Ind, Mel, Phil, Vie), taro (Mel) and maize (Ind). Green revolution dynamics for each geopolitical entity are reflected by time paths for paddy rice. For each trendline, the intercept is reflective of past technological change in a particular cropping system.

Over 1961-2016, rice yield growth was surpassed by that of other (BC-reliant) commodities at different times. In Melanesia, yield growth of sugarcane (158%) and coconut (106%) amply surpassed that of rice by 1962 and 1964, respectively. In Indonesia, rice yield growth was consistently higher than that of other commodities until 1989 and 1995 when it was surpassed by that of banana and maize respectively. In the Philippines, yield growth in BC-reliant banana amply surpassed that of rice during 1974-2004. In Thailand and Vietnam, yield gain in coconut was most pronounced over 1998-2010, a period that coincided with BC interventions against the leaf-feeding *B. longissima* (Nakamura et al., 2006).

## Discussion

Biodiversity-driven ecosystem services such as biological control (BC) underpin sustainable food systems and safeguard societal wellbeing in the face of environmental change (Bale et al., 2007; Dainese et al., 2019). As a key constituent of global change, pest invasions result from an ecological imbalance that occurs when non-native species are released from BC-mediated regulation (Elton, 1958; Keane & Crawley, 2002). Scientifically-guided BC restores such balance, reconstitutes ecosystem service delivery and thereby alleviates pest-induced economic, environmental and societal impacts (Hoddle, 2004). Here, we illuminate how 76 different insect BC agents (equaling at least 175 unique country-level BC interventions) mitigated min. 45 pest targets over 1918-2018. BC delivered durable pest control in myriad Asia-Pacific agriculture and livestock production sectors, permitting an associated yield-loss recovery up to 73%, 81% or 100% in crops such as cassava, banana or coconut (Waterhouse et al., 1998; Thancharoen et al., 2018). The ensuing economic dividends are substantial, with pest-induced losses amounting to US $6.7, $4.3 and $8.2 billion annually for the above crops (Fig. 3). These annually-accruing benefits compare favorably with the one-off US$4.5 billion Asia-wide benefit of improved rice germplasm (Raitzer & Kelley, 2008). By remediating persistent pest shocks, BC reasonably promoted technological change and triggered growth within several agricultural sub-sectors. Nations such as Indonesia that purposely prioritized BC enjoyed productivity gains in pest-afflicted banana (1,222%), maize (572%) and cassava (322%), far surpassing those of paddy rice (303%) (Fig. 5). Moreover, direct on-farm monetary benefits of BC equal 8.6-14.7% current GDP of several Pacific island states and 0.5% GDP of industrialized economies such as Indonesia. Our work constitutes the first empirical demonstration of how BC solidified the agrarian foundation and lifted overall performance of various Asia-Pacific economies (Huffaker & Caltagirone, 1986). Doing so places BC on equal footing with other biological innovations such as GR-enhanced germplasm, spotlights its transformative impacts, and celebrates the century-long achievements of dedicated yet often unacclaimed insect explorers and biological control pioneers.

Technological change and its associated productivity gain are engines of agricultural growth (Ruttan, 2002), yet aren’t exclusively due to improved genetics, mechanization or chemical technologies but also relate to agro-ecological measures (de Janvry, 2010; van der Ploeg et al., 2019). Invasive pests represent large productivity shocks that can debilitate a country’s agriculture sector and slow economic growth (Haggblade et al., 2007), with their impacts likely inflated in small economies (Cardia, 1991) or when degrading ‘point-source’ resources e.g., oilseed palm plantations and the associated export-geared value chains (Lederman & Maloney, 2008). BC-induced suppression of invasive pests removes bottlenecks in farming operations, enabling permanent shifts of agricultural production isoquants (Ruttan & Hayami, 1998) and feeding forward into the overall economy (de Janvry, 2010; Wiggins et al., 2010). Strong linkages exist between agricultural growth and the rural non-farm economy, with respective growth multipliers ranging between 1.6 and 1.8 for Asia (Haggblade et al., 2007). Considering how the multipliers of GR-induced agricultural growth varied between resource-endowed and marginal settings (Pingali, 2012; DeFries & Nagendra, 2017), we argue that BC in (smallholder) non-rice crops such as cassava, banana or maize enabled a stable, equitable ‘miracle’ growth of economies such as Indonesia (Timmer, 1998; Dunning, 2005). In comparison with GR seed x fertilizer technologies, BC acted as a self-sustaining innovation across sites with variable endowment and lifted productivity irrespective of local socio-economic or biophysical context (Ruttan & Hayami, 1998). Furthermore, as a cost-free alternative to input-intensive measures for pest control, BC improved financial solvency of farm operations - thereby alleviating poverty (Wiggins et al., 2010) and bolstering demand for consumer goods (Haggblade et al., 2007). We hypothesize that BC elevated the purchasing power of farm households while suspending backward linkages to (foreign-owned) agrochemical suppliers (Vogel, 1994; Folke et al., 2019), with the ensuing multiplier effect to domestic economies conceivably being higher than GR technology packages. These household-level savings can be substantial, as exemplified by the US$ 1.3-2.3 billion yearly insecticide expenditure to tackle the invasive *Plutella xylostella* (Linnaeus) in the world’s cabbage crops (Zalucki et al., 2012). Lastly, we theorize that a sequence of BC successes in e.g., Indonesia or Philippines induced social-ecological change (Ruttan & Hayami, 1998; Yletyinen et al., 2019) that ultimately facilitated the roll-out of effective input-reduction initiatives such as UN-FAO’s Farmer Field School program (Bottrell & Schoenly, 2012; Thorburn, 2015).

Our projected US$ 22.6 billion regional benefit conceivably represents only a fraction of the full economic weight of biological control. Several caveats exist in our analysis, e.g., uneven reporting of BC outcomes between countries, a tendency to overstate impacts, irregular congruence between initial assessments and subsequent ground-truthing (Cock et al., 2016) and fragmented knowledge of in-country variability in incidence or yield impact of invaders. Conversely, production statistics for multiple, pest-afflicted commodities are incomplete, unreliable or absent (Suppl. Fig 2). Also, our estimates do not account for cosmetic damage and quality loss to farm produce, loss of export markets, pest management costs (Zalucki et al., 2012), price shocks or rippling effects either backward or forward (i.e., processors of raw produce such as copra) along the value chain – with the latter amplified by trade (Reardon, 2015; Wyckhuys et al., 2018). Moreover, various BC-reliant crops (e.g., bananas, root crops, breadfruit) are not extensively traded in the marketplace and irregularly assigned a monetary value (Waterhouse et al., 1998) while crops such as cassava overcompensate low-level pest attack (Thancharoen et al., 2018). The 30% lower estimated value of *E. thrax* biological control in PNG, as compared to full-fledged assessments (Waterhouse et al., 1998), exemplifies the conservative nature of our calculations.

The discipline of biological control finds itself at a crossroads. As BC resilience is directly tied to global environmental change (Martin et al., 2019), habitat loss, agrochemical pollution and the insect biodiversity decline are causing net negative impacts (Geiger et al., 2010; Rusch et al., 2016; Dainese et al., 2019). Though BC can help resolve multiple sustainability challenges and stem a global proliferation of pesticide use (Bernhardt et al., 2017), it suffers from a steady decline in institutional capacity, lacking public recognition (Warner et al., 2011; Barratt et al., 2018) and stringent regulations (Heimpel & Cock, 2018). However, modern BC explicitly balances ecological risks with multi-faceted benefits, and its judicious use can thus safely defuse invasive species impacts (Bradshaw et al., 2016; Paini et al., 2016), ease vector-borne disease burden (Benelli et al., 2016), and exert stabilizing effects on global commodity markets (Wyckhuys et al., 2018). Moreover, as BC resolved 80% losses of breadfruit i.e., the ‘staff of life’ of Pacific Islanders (Kirch & Ralu, 2007) or enhanced dietary carbohydrate intake of root, tuber and banana-consuming communities in eastern Indonesia and PNG, it raised the carrying capacity of local food systems (Mottesharrei et al., 2014; Dangour et al., 2017; Gillespie & van den Bold, 2017), enhanced workforce productivity (Haggblade et al., 2007) and generated ‘win-win’ outcomes for human and natural capital (Rasmussen et al., 2018).

The Asia-Pacific region comprises multiple prime biodiversity (island) ‘hotspots’ with high rates of endemism (Bellard et al., 2017), which prove highly susceptible to species invasion (Reaser et al., 2007) or to misguided BC releases e.g., as conducted pre-1945 in certain US territories (Henneman & Memmott, 2001). Given that modern BC aims to conserve local biota and restore native ecosystems (van Driesche et al., 2010), ample benefits can be reaped from collaboration between conservationists and crop protection scientists. Such trans-disciplinary work e.g., in Asia-Pacific island ‘laboratories’ can define common denominators of success and raise rates of natural enemy establishment (38%), BC success and ecological safety (DeBach, 1962; Stiling & Cornelissen, 2005). The chronicled BC events were made possible through dedicated science, globe-spanning partnerships and sustained funding commitments. As today’s agricultural R&D investments dwindle (Alston & Pardey, 2014), the relatively modest budget needs for BC should be contrasted with its ample spillover benefits (Naranjo et al., 2015), the 40% of public funds diverted to ‘maintenance research’ e.g., to resolve pesticide-induced herbivore resistance (Sparger et al., 2013; Jorgensen et al., 2018) and the vast sums of money that are currently re-routed to computational or molecular sciences (Warner et al., 2011).

The Green Revolution is rightly credited with averting Malthusian famines, spurring structural transformation and delivering vast economic dividends (Threwavas, 2002; Pingali, 2012; Gollin et al., 2018). While ‘packaged’ GR seed x chemical technologies in Asian paddy rice systems did secure an elevated, stable supply of energy-dense foods, we argue that these achievements were paralleled by biodiversity-based innovations. In GR-beneficiary nations such as Indonesia or Philippines, successive waves of invasive species impacted 8-13 key agricultural commodities - often triggering sudden food shortages, crippling rural agro-industries or cascading into chronic societal crises (Waterhouse et al., 1998; Ricciardi et al., 2011; Tittonell & Giller, 2013; Yletyinen et al, 2019). Scientifically-guided BC across this diverse portfolio of commodities solidified the agricultural base of heterogeneous rural settings, lowered poverty vulnerability and ultimately enabled stable, sustainable and equitable growth of agricultural and non-farm sectors (Weinberger & Lumpkin, 2007; de Janvry, 2010). Moreover, the annually-accruing socio-economic and environmental spin-offs of 76 carefully-selected BC agents surely outpace those of input-intensive GR technologies (Bernhardt et al., 2017; Springmann et al., 2018). Now, as a new dawn is breaking for global agriculture (de Janvry, 2010; Pretty et al., 2018; Eyhorn et al., 2019), our work highlights the transformative potential of biodiversity-based innovations to secure a profitable production of sufficient, safe and nutrient-dense farm produce, while preserving our common natural heritage.

## Acknowledgements

We are grateful to Frances Williams for valuable comments that improved an earlier draft of the manuscript. The development of this manuscript and its underlying research were partially funded through the Australian Centre for International Agricultural Research (ACIAR) HORT/2016/185. The maintenance of BIOCAT and MC’s inputs were supported by the CABI Development Fund (supported by contributions from ACIAR, the UK Department for International Development, the Swiss Agency for Development and Cooperation and others). CABI is an international intergovernmental organisation and gratefully acknowledges the core financial support from its member countries; see https://www.cabi.org/about-cabi/who-we-work-with/key-donors for details.

## Author contributions

KAGW conceived and designed the experimental approach; KAGW performed trials and collected the data; KAGW and WWZ analyzed the data; all authors co-wrote the paper.

## Data Availability

All data underlying the analyses are made available through Dryad Digital Repository at XXX.

## Materials & Methods

### Data collection

As the basis for our assessment of historical biological control (BC) efforts, we consulted the BIOCAT2010.2 database (Greathead & Greathead, 1992; Cock et al., 2016). More specifically, we queried its 2017 updated version and extracted country-level records on natural enemy identity, release date, target pest and crop, and fate of the resulting biological control endeavor for all geopolitical entities within Southeast Asia (including South China), Melanesia, Micronesia and Polynesia. Only records of successful natural enemy establishment were retained, and BIOCAT classifications were used to record locality-specific impacts of a given BC event. On the latter, we exclusively considered BC events that resulted in either full or partial control of the target pest in any of its geopolitical areas. Detailed data on historic biological control interventions spanning an entire century (1918-2018) were thus collated, from which records for the US State of Hawaii were deleted. Similarly, for subsequent economic analyses, we removed data from the US territories of Guam and American Samoa and solely retained interventions that were aimed at resolving pest issues in the agriculture and livestock production sector.

For macro-economic interpretations, the above dataset was complemented with a non-exhaustive review of the global literature for specific information pertaining to each invasive pest and introduced natural enemy. Google Scholar (GS) was used as a search engine to extract relevant records from the scientific literature, running queries between May 15 and August 20, 2019. More specifically, we screened the literature for data on pest-induced yield loss, pest incidence, natural enemy establishment and geographical spread, predation or parasitism levels, and extent of pest population reduction as mediated by biological control. For each literature record, information was extracted on geographical locality and host plant association of a given pest or natural enemy. If relevant data were available from publications that emanated from within the Asia-Pacific region, these were prioritized for subsequent analyses and economic interpretations. If local data were not available for certain metrics, these were complemented with literature records from other areas in the invaded range of a given pest (and introduced natural enemy). Despite eventual shortcomings within the original field studies, we tried to gauge pest-induced crop losses and BC-mediated yield recovery in a comprehensive fashion (see below). The latter was done using the BIOCAT classifications for BC success rate, lumping ‘complete’ and ‘substantial’ control and treating ‘partial’ success separately.

For each impacted crop, country-specific information on production statistics and pricing records was extracted from FAOSTAT (http://www.fao.org/faostat/). Crop- and country-specific data were thus collated on production level (harvested area, ha), yield (t/ha), total production quantity (tonnes) and producer price (US$) for 2015, the most recent year for which data from most countries were available. As such, we use the most complete and reliable production statistics, but do not account for certain commodities that experienced important historic reductions in cropping area (e.g., cassava in Indonesia). For geopolitical areas where no local information was available on producer prices for a given commodity, regionally-averaged prices were used.

### Data analysis and interpretation

Upon consolidation of all datasets, we plotted the number of BC introductions over time and listed the identity of natural enemies (and associated pest targets) that were released in each geopolitical area. BIOCAT records were used to compute both country-level and regional rates of natural enemy establishment and success of ensuing BC efforts. BC success rates were further compared between taxonomic hierarchies (i.e., order, species).

To quantify maximum potential losses due to an invasive pest in a given area, we multiplied the maximum pest-inflicted productivity loss (i.e., percentual yield reduction, as extracted from the global literature) on its main host crops with their respective 2015 local production quantity and producer price. Doing so yielded an estimate of maximum yearly pest-inflicted economic losses, computed for current production scenarios in a particular area. For host crops of which pest-induced productivity loss was unknown, a default value of 25% yield reduction was used. This rate is comparable to documented invasive pest impacts in the Asia-Pacific region and deemed to be conservative as no further (off-farm) economic impacts were accounted for. As such, we quantified the total economic loss as incurred by local farmers or livestock producers. This method did not account for eventual further economic losses along the value chain for a given crop or livestock item (including lowered consumer prices due to compromised produce quality), farm-level expenditures for invasive pest management or foregone export earnings due to e.g., restricted international market access. Also, by using present-day FAOSTAT producer prices, we disregarded (often substantial) price elasticity in local markets following pest-induced yield and production shocks. To estimate the economic benefits of a BC intervention within a certain geographical setting, we multiplied the respective (pest-induced) economic loss with a BIOCAT-derived metric reflecting in-country success of the BC effort. For natural enemies that resulted in variable rates of BC success across its introduced range, we used the highest level of success reported. For fully- or partially-successful BC interventions, we considered a respective 100% or 50% yield loss recovery. As such, we are prone to capture an upper-bound of BC-mediated productivity gain, but concurrently do not account for monetary benefits that accrue through lowered pest management costs, restored product quality, price shocks or cascading impacts along the produce value chain (e.g., Zalucki et al., 2012). Both pest-induced economic losses and BC-related benefits were expressed in absolute terms and as percentage of the 2015 national gross domestic product (GDP). To allow for eventual inter-country comparisons, GDP figures were calculated at purchasing power parity (PPP).

### Econometric analyses

To track year-by-year land and labor productivity growth for each of the Asia-Pacific geopolitical entities, we employed the graphical technique of Hayami & Ruttan (1971). More specifically, we expressed the value of country-level agricultural production and plotted the respective labor productivity as a measure of land productivity. Aggregate agricultural production figures were drawn from FAOSTAT, with production value expressed in 2004-2006 average purchasing power parity (PPP) prices (i.e., international dollars). Estimates of agricultural labor employment and total agriculture area were obtained from the World Bank Data portal (https://data.worldbank.org/). This allowed visualizing starting points and temporal patterns in productivity growth for each country over a 1960-2016 window, though agricultural employment figures were only available for most countries from 1991 onward.

Next, we assessed the relative contribution of historic BC interventions to country-level land productivity growth over 1960-2016 in three different ways. First, to gauge the relative impact of GR rice technologies on growth trends within specific agricultural sub-sectors, we contrasted value of agricultural output of the locally-prevailing food staple (i.e., paddy rice or taro + sweet potato) with that of aggregate agricultural output for a specific country or geopolitical entity (e.g., Melanesia). Second, we plotted growth trajectories of the above agricultural sub-sectors and measured inter-decadal proportional change in respective output, as compared with a 1961-1970 baseline. Third, for selected geopolitical entities, we drew time-paths for different (BC-affected) agricultural commodities including paddy rice by plotting inter-annual percentual growth in area (hectare) versus yield (tonne/hectare) against a 1961 base year. This allowed a quantitative partitioning of historical output growth for specific commodities among area expansion and yield enhancement, thus reflecting relative magnitude and type (i.e., land- or labor-saving) of commodity-specific technological change. For BC-impacted commodities, we equaled compared year-by-year changes in yield growth with those of the locally-prevailing food staple (i.e., paddy rice or taro – for Melanesia).

